# Mechanically stable knot formed by strand threading in Zika virus RNA confers RNase resistance

**DOI:** 10.1101/2020.07.01.183129

**Authors:** Meng Zhao, Michael T. Woodside

## Abstract

Exoribonuclease-resistant RNAs (xrRNAs) from viruses prevent digestion by host exoribonucleases, creating sub-genomic viral RNAs that can enhance infection and pathogenicity. Novel knotted structures in xrRNAs are proposed to act as mechanical road-blocks to RNases. Studying an xrRNA from Zika virus with optical tweezers, we found that it was the most mechanically stable RNA structure yet observed. The knot folded by threading the 5′ end into the cleft of a Mg^2+^-coordinated three-helix junction before pseudoknot interactions closed a ring around it. Both the threading and pseudoknot were required to generate the extremely force-resistant knot, whose formation correlated directly with RNase resistance both in the wild-type xrRNA and a low-resistance mutant. This work clarifies the folding and mechanism of action of an important new class of RNA.

---

Cells infected by viruses typically attempt to digest the viral RNA with RNase enzymes. By inserting structures that are resistant to RNase digestion into the viral genome, viruses can turn this defense back against the host cell, using host RNases to generate sub-genomic RNAs in downstream protected regions. Such sub-genomic fragments have been found to interact with host proteins and interfere with multiple cellular processes, enhancing viral transmission and pathogenicity (*1-5*). First discovered in flaviviruses (*6, 7*), exoribonuclease-resistant RNAs (xrRNAs) capable of resisting degradation by 5′→3′ exoribonucleases without cofactors have since been found in other virus families (*8-12*). They may therefore present a fruitful new target for anti-viral therapeutics.

Structural studies of xrRNAs (*11-14*) revealed a previously unknown fold architecture featuring a single strand threaded through a ring closed by secondary and tertiary interactions to form a knot-like structure, which we term a ring-knot. The ring-knot fold is proposed to be responsible for exoribonuclease resistance (XR) by blocking digestion mechanically, preventing the unfolding of xr-RNAs by the helicase activity of RNases (*15*). This hypothesis remains untested, however, because the mechanical stability of xrRNAs has not yet been measured. Furthermore, although single-molecule fluorescence measurements have probed the dynamics of the ring-closing pseudoknot (*11*), the full sequence of events leading to the formation of the ring-knot—placing the folding of secondary-structure elements and the formation of the tertiary contacts in their relevant contexts—has yet to be established. As a result, it remains unclear how the knot folds, which interactions are critical to the putative mechanical resistance, and how these interactions relate to XR.

To address these questions, we quantified the resistance to mechanical unfolding of an xrRNA from Zika virus using single-molecule force spectroscopy (*16*), applying an increasing force to the ends of a single Zika xrRNA molecule with optical tweezers (Fig. 1A) until the xrRNA unfolded, and then reducing the force to watch it refold. This approach allows the complete folding process, starting from the fully unfolded state, to be observed directly. Moreover, by measuring the extension of the molecule as its structure changes, the number of nucleotides involved in each transition can be determined and the structure of intermediate states deduced. RNA containing the xrRNA sequence (Fig. 1B, Table S1) flanked on either side by kilobase-long handle sequences was annealed to complementary single-stranded DNA handles attached to beads held in optical traps (Fig. 1A). The traps were then moved apart to ramp up the force on the RNA before being brought back together to ramp the force down, waiting 0.5–5.5 s at near-zero force for the xrRNA to refold before pulling again. This procedure generated force-extension curves (FECs) reflecting the structural changes during unfolding and refolding: unfolding (refolding) of substructures within the RNA was detected as characteristic “rips” in the curves (Fig. 1C–F), where the extension abruptly increased (decreased) concomitant with a decrease (increase) in the force.

**Fig. 1:**
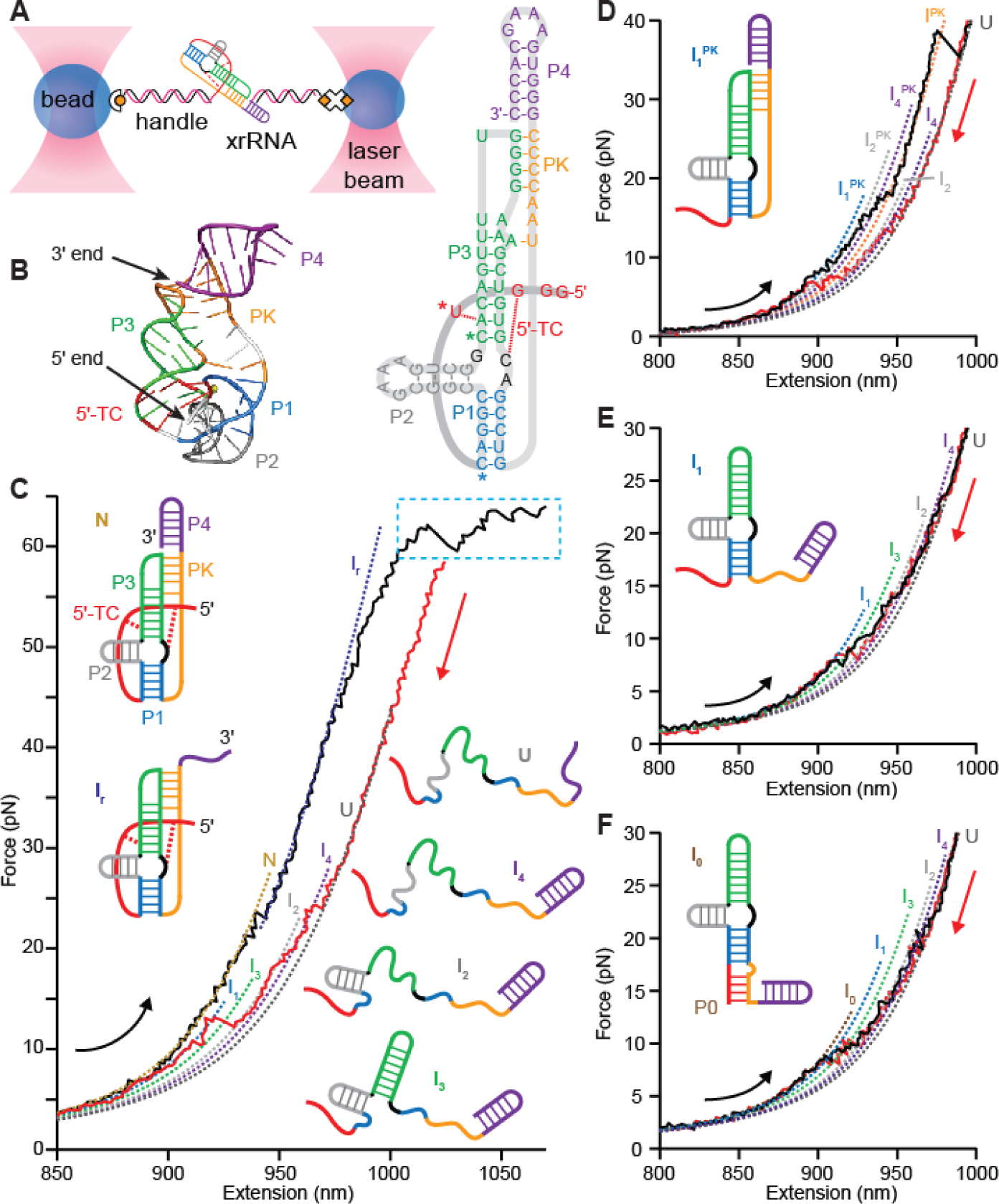
Single-molecule force spectroscopy of Zika xrRNA. (**A**) A single xrRNA is held under tension via DNA handles bound to beads held in optical traps. (**B**) Left: structure of Zika xrRNA showing color-coded helices (P1–P4), pseudoknot (PK), and threading contacts (5′-TC). Right: xrRNA sequence. (**C**) Most unfolding FECs (black) don’t show complete unfolding until the overstretching transition (cyan); refolding FECs (red) show sequential formation of helices. Dashed lines: WLC fits for each state (native N, intermediates I_x_, unfolded U). Insets: structural cartoons for each state, color-coded as in (A). (**D**) Roughly 4% of FECs show unfolding of an unthreaded pseudoknot structure (inset) at lower force, ∼30–50 pN. (**E**) Roughly 15% of FECs involve secondary structure only (inset), with helices P1–P4 unfolding at forces of ∼10–20 pN. (**F**) Roughly 1% of FECs involve a secondary structure only but with a non-native extension of helix P1 (P0, inset).

Examining the unfolding of the wild-type xrRNA in the presence of 4 mM Mg^2+^, the great majority of FECs (∼80%) showed a small rip at ∼20 pN, but then typically remained featureless up to ∼60 pN (Fig. 1C, black). Fitting the FECs before and after the low-force rip to extensible worm-like chain (WLC) models (Fig 1C, dotted lines) (*17*), we found a contour-length change of Δ*L*_c_ = 6.1 ± 0.4 nm, only a small fraction of the 38.3 nm expected for complete unfolding (*14*). The xrRNA thus remained in a mostly folded intermediate, denoted I_r_, even at 60 pN—well above the unfolding forces reported for any other RNA structures (*18-24*) and in the same range as the duplex over-stretching transition (*25*). Indeed, in most of these curves, unfolding the xrRNA completely required holding it at the overstretching force for at least 2 s; as a result, I_r_ unfolding occurred at the same time as handle overstretching and was difficult to distinguish (Fig. 1C, cyan, Fig. S1A). After waiting 2 s at ∼65 pN, however, refolding curves displayed the full Δ*L*_c_ expected (Fig. 1C, red, Fig. S1B), revealing multiple intermediates that were well fit by WLC models (Fig. 1C, dashed lines) yielding the contour-length changes for each state (Table S2).

Turning to the unfolding FECs without extreme mechanical resistance (∼20% of all FECs), roughly 1/5 (4% of all FECs) unfolded through 3 intermediate states at low forces (Fig. 1D, Fig. S1C), in the 10–20 pN range characteristic of secondary structures (*26*), followed by a final transition at higher force, in the ∼30–50 pN range characteristic for unfolding tertiary structures (*18-22*). The total Δ*L*_c_ for complete unfolding in these FECs, 36 ± 1 nm, was slightly shorter than for the native fold, indicating that the xrRNA started in a state close to the native fold. The remaining FECs without extreme mechanical resistance (16% of all FECs), in contrast, unfolded through low-force intermediates only, indicating that no tertiary contacts had formed. Most (15% of all FECs) had a total Δ*L*_c_ of 31.3 ± 0.8 nm (Fig. 1E, Fig. S1D), shorter than expected and hence indicating the xrRNA was not fully folded, whereas a very few (1% of all FECs) had an additional low-force transition resulting in a total Δ*L*_c_ of 36.9 ± 0.5 nm (Fig. 1F, Fig. S1E).

To interpret these results, we compared the Δ*L*_c_ values from WLC fits to the values expected for unfolding various substructures of the native fold (Fig. S2, Table S2). For the FECs with high mechanical resistance (Fig. 1C, Fig. S1A), the low-force rip corresponded to unfolding helix P4 (Fig. 1B, purple), leading to the structure for I_r_ illustrated in Fig. 1C. The refolding preceding such unfolding curves was consistent with sequential formation of first P4 (producing state I_4_), then P2 (producing I_2_), P3 (producing I_3_), and finally P1 (producing I_1_); the pseudoknot (denoted PK) and 5′-end tertiary contacts (denoted 5′-TC) responsible for high unfolding forces must have formed subsequently at too low a force to observe reliably. For the FECs without high-force unfolding events, the unfolding for the majority (Fig. 1E, Fig. S1D) was consistent with starting in state I_1_ containing native secondary structure only (Fig. 1E, inset) and proceeding through I_3_, I_2_, and I_4_ to the unfolded state (U); the unfolding of the others (Fig. 1F, Fig. S1E) was consistent with starting in state I_0_ incorporating a non-native helix P0 (Fig. 1F, inset) and then proceeding through the same sequence of intermediate states to U. Finally, for the remaining 4% of FECs (Fig. 1D, Fig. S1C), the unfolding matched the lengths expected for a non-native state (Fig. 1D, inset) with all helices formed and the ring closed by PK but with the 5′ end unthreaded through the ring: first P1 was unfolded, then P2, P4, and finally (at high force) PK. The refolding curves preceding these unfolding FECs were very similar to those seen in the other cases (Fig. S3), forming in sequence I_4_, I_2_, I_3_, and finally a variant of I_1_ with the pseudoknot interaction intact (denoted I_1_^PK^).

The picture revealed by these measurements is intuitive: refolding involves forming each helix, in the order P4, P2, P3, and P1, followed by one of four options: (i) threading of the 5′ end before PK closure to achieve the full native tertiary structure (N, 80% likelihood); (ii) closure of PK before threading the 5′ end to achieve only partial tertiary structure (I_1_^PK^, 4% likelihood); (iii) neither threading nor PK closure, leaving an unconstrained 5′ end (I_1_, 15% likelihood); or (iv) alternative secondary structure formation inhibiting 5′-end threading (I_0_, 1% likelihood). The difference in threading between N and I_1_^PK^ makes it impossible for them to inter-convert without unfolding the pseudoknot. The proportion of threaded and unthreaded states did not change significantly when the waiting time between successive pulls at near-zero force was reduced from 5.5 s to 0.5 s (Fig. S4), indicating that threading and PK closure occurred rapidly.

To confirm this interpretation, we re-measured the FECs in the presence of anti-sense oligonucleotides complementary to specific regions of the sequence, to prevent formation of the targeted structures and interactions (Fig. 2A). Blocking P4 from forming with oligo 1 abolished the rip at ∼20 pN attributed to P4 unfolding but left the resistant intermediate I_r_ intact, as expected (Fig. 2B(i)). In fact, the fraction of curves containing I_r_ increased slightly to ∼90% using oligo 1, while ∼10% of FECs started in a state (denoted I_1b_) with P1–P3 formed but no tertiary contacts (Fig. S5), suggesting that steric hindrance from P4 may hinder 5′-end threading under normal conditions. Blocking both P4 and PK with oligo 2, in contrast, abolished I_r_ entirely (Fig. 2B(ii)); FECs instead unfolded from state I_5b_ (P1–P3 and 5′-TC but no PK), in the force range ∼25–55 pN, or from state I_1b_, at ∼20 pN (Fig. S6), indicating that the pseudoknot is essential for the extreme mechanical resistance of the xrRNA. These results show that 5′-TC can form independently of PK, and that the intermediate I_5_ (Fig. S2A) with helices and 5′-TC formed but not PK—which is necessary for threading the 5′ end before loop closure but not observed directly in the absence of anti-sense oligos—is indeed metastable. Lastly, blocking P1 and 5′-TC with oligo 3 abolished all tertiary structure (Figs. 2B(iii), Fig. S7), indicating that native pseudoknot formation requires P1. Revisiting the FECs without oligos but removing Mg^2+^, which is required for native 5′-TC (*14*), most FECs (∼67%) revealed folding of only native secondary structure, similar to the oligo 3 results (Fig. S8). The remaining FECs showed unfolding lengths and forces consistent with formation of a variant pseudoknot interaction (denoted PK′) without 5′-end threading (Fig. 2B(iv), Fig. S8), suggesting pseudoknots can form independently of 5′-TC. The length changes associated with the structural transitions in all FECs with oligos and without Mg^2+^ are listed in Table S3.

**Fig. 2:**
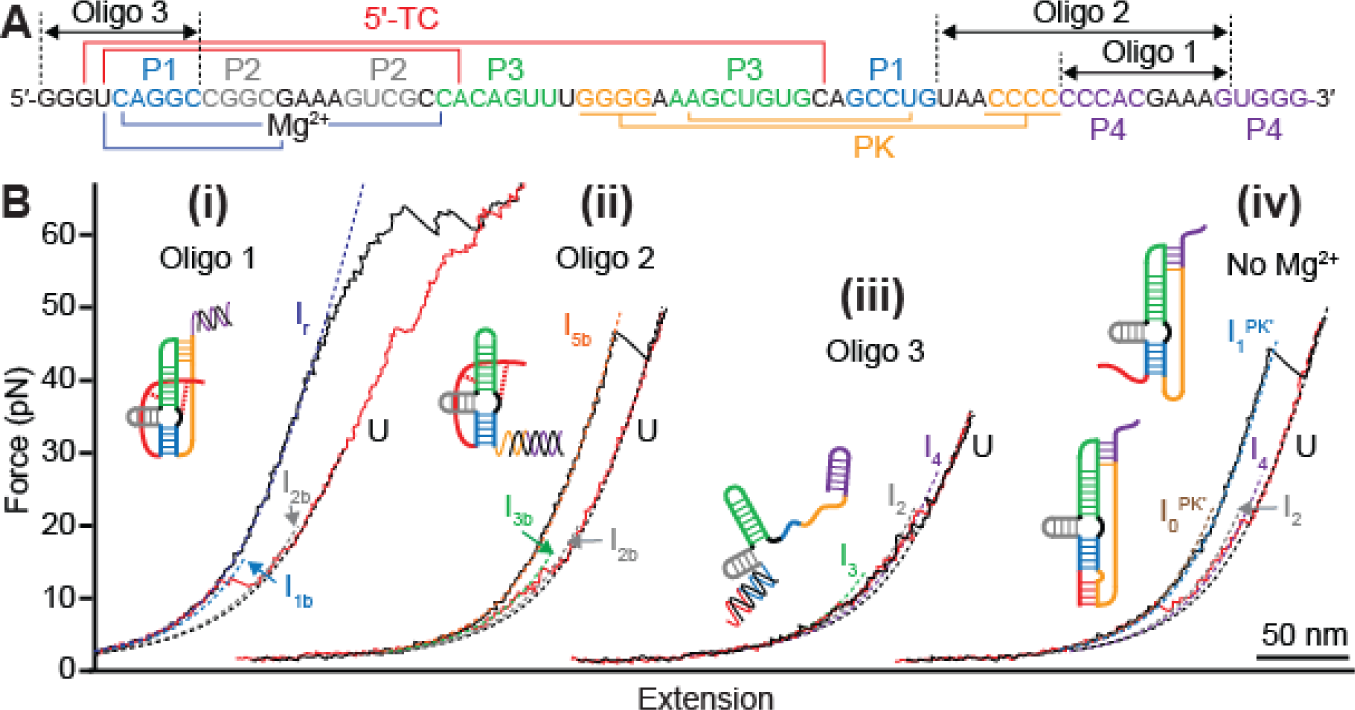
FECs in presence of anti-sense oligos. (**A**) Locations where anti-sense oligos pair with xrRNA sequence: oligo 1 blocks formation of P4, oligo 2 blocks P4 and PK, oligo 3 blocks P1 and 5′-TC. (**B**) Representative unfolding (black) and refolding (red) FECs with (i) oligo 1, (ii) oligo 2, (iii) oligo 3, and (iv) in the absence of Mg^2+^. Insets show structural cartoons of the state in which unfolding FECs start. Dashed lines: WLC fits.

Combining all of these results provides a comprehensive view of the unfolding and refolding pathways (Fig. 3) and suggests the roles played by different parts of the xrRNA. The critical state responsible for the unprecedented mechanical stability of the xrRNA is the intermediate I_r_, not the full native state: P4 can unfold without compromising the mechanical strength of the knot. Intriguingly, mutational studies suggest that P4 is not essential to XR activity, either (*13, 14*). Instead, P4 seems to play an assisting role as a “buckle”, helping to keep the ring compact and prepare the way for native 5′-TC and PK interactions by preventing the formation of the more extended non-native pseudoknots (like PK′) seen when P4 does not fold, which do not lead to threading (Fig. S8). The other crucial state is I_1_, where the folding pathway divides depending on whether PK or 5′-TC forms first, since each can form independently of the other. If PK forms before the 5′ end threads into the cleft in the 3-helix junction formed by P1–P3, then no threading is possible, and the RNA is trapped topologically in a less mechanically stable state. Only if the Mg^2+^-mediated 5′-end threading occurs before PK closure is the native state with exceptional mechanical resistance achieved.

**Fig. 3.**
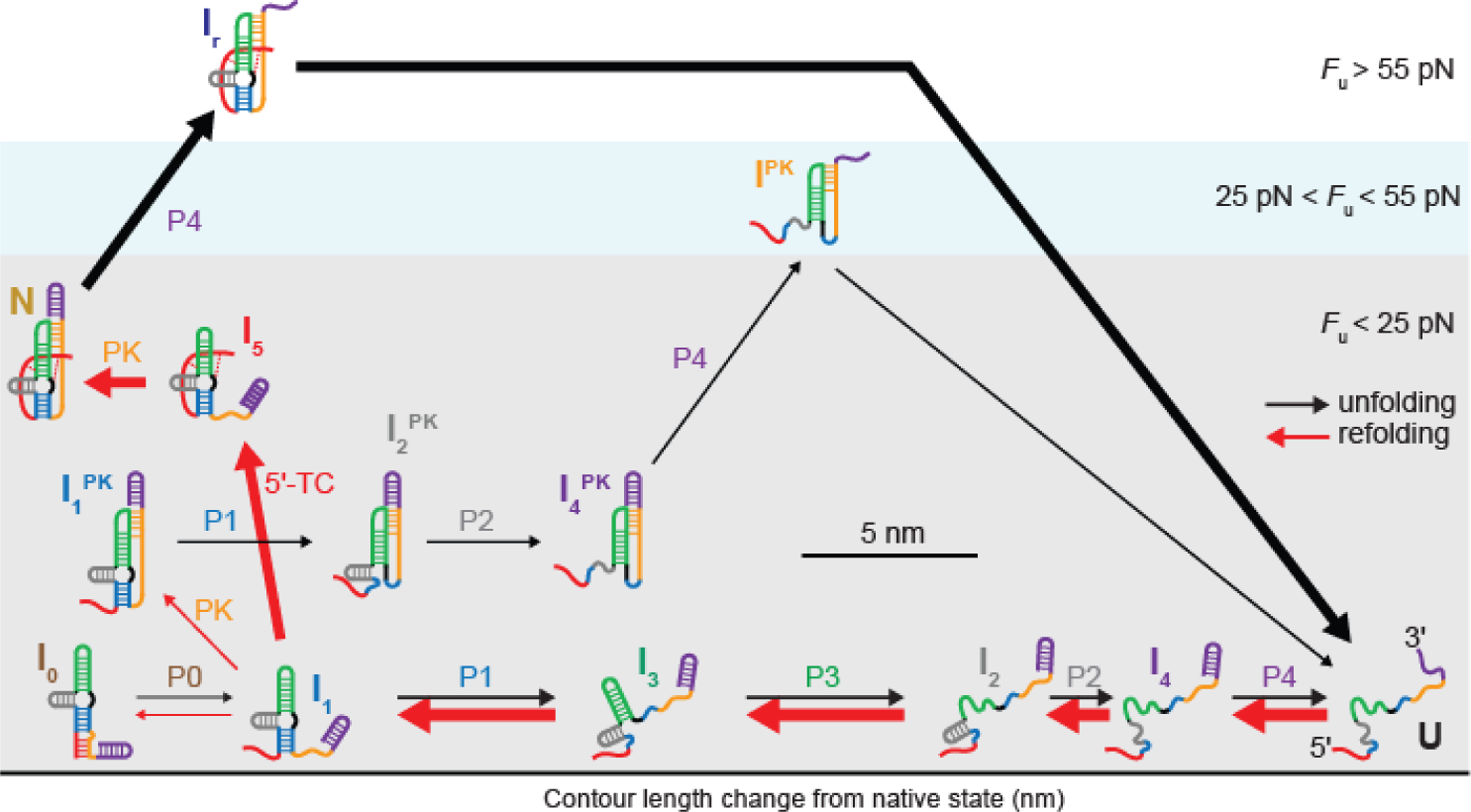
Unfolding and refolding pathways. The position of each state (shown as a cartoon of its structure) reflects its unfolding force and Δ*L*_c_ relative to N. Arrows (black: unfolding, red: refolding) denote transitions, where the indicated structural element unfolds/refolds; their widths are proportional to the fraction of FECs in which the given transition occurs.

Finally, to test if the unusual mechanical properties of I_r_ underlie XR activity, we studied the effects of mutating base C22 to G, which reduces XR activity significantly—80 ± 4% vs. 9 ± 5%—putatively by replacing the 5′-TC involving G3 and C44 with the base-pair G22:C44, lengthening P3 by 1 base-pair (*13, 14*). However, only 32% of unfolding FECs were consistent with this scenario of a longer P3 and shorter P2: 15% containing secondary structure only (a variant of I_1_) and 17% also featuring PK′ (Fig. 4A, Fig. S9). The majority, 68%, were consistent with the wild-type helix P3 and instead had P2 lengthened by a C9:G22 base-pair (Fig. 4B, Fig. S9, Table S4), leading to an increased unfolding force (Fig. 4C), whereas P1 was shortened by 1 base-pair. Most of these curves (53% of all FECs) again showed only secondary structure (another variant of I_1_), but a significant minority (15% of all FECs) retained a ring-knot architecture despite the distortions to P1 and P2, displaying an extremely high-force intermediate after P4 unfolding analogous to I_r_, denoted I_r_^m^. I_r_^m^ can form because C44 remains unpaired, allowing for full 5′-TC formation (Fig. S10). However, the mutation disfavors this configuration, resulting in a much-reduced incidence of the ring-knot.

**Fig. 4:**
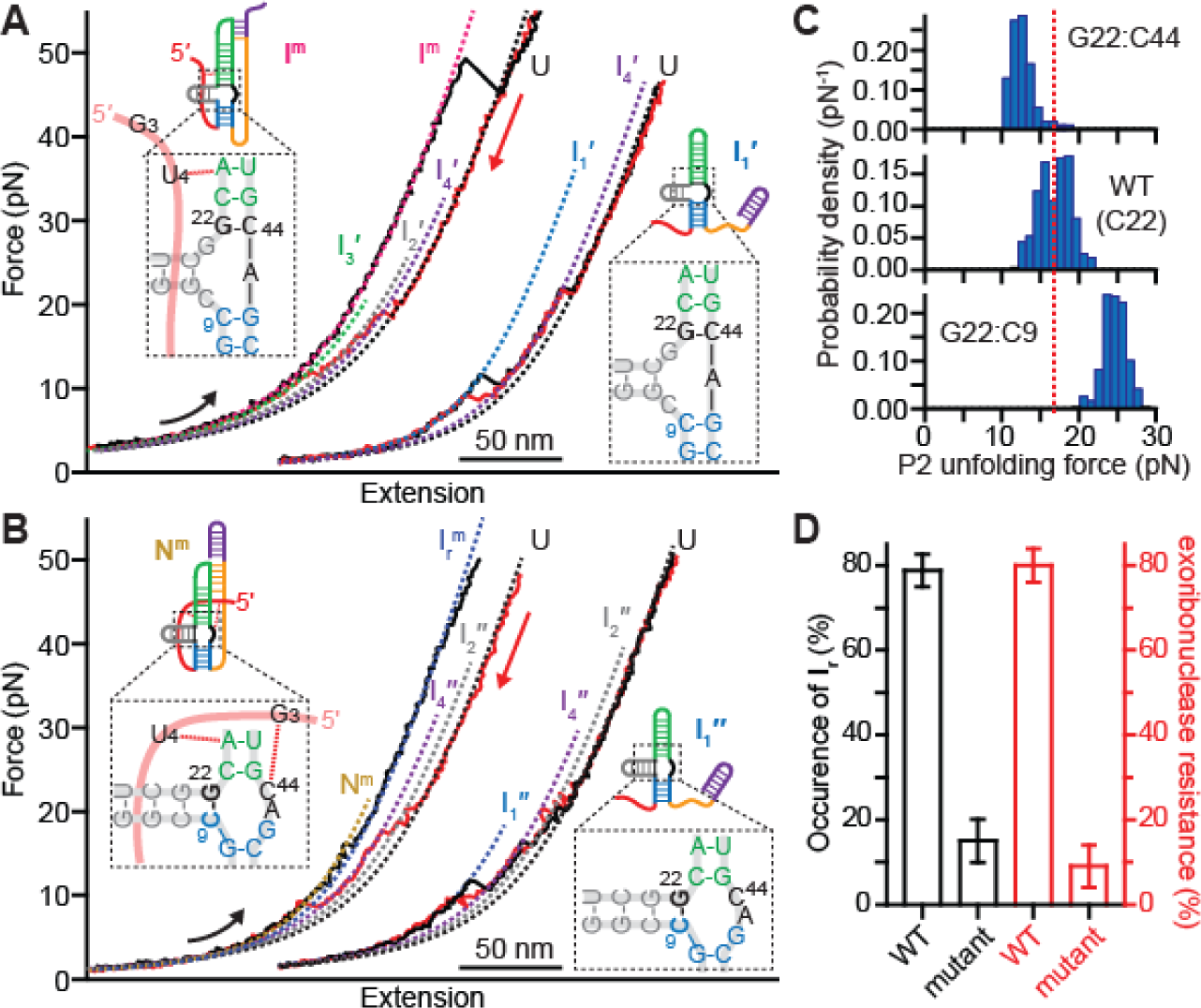
Folding of C22G mutant. **(A)** 1/3 of unfolding (black) and refolding (red) FECs were consistent with G22:C44 base-pairing, which prevents 5′-end threading: 17% (left) featured PK′ (non-native pseudoknot), 15% (right) helices only. Insets: cartoons of structures at start of unfolding FECs. (**B**) The remaining FECs were consistent with G22:C9 base-pairing: 53% (right) featured helices only, but 15% (left) featured a fully threaded ring-knot. Insets: cartoons of structures at start of unfolding FECs. (**C**) With G22:C44 pairing, P2 is shorter than in wild-type and has a lower unfolding force (top), whereas with G22:C9, P2 is longer and has a higher unfolding force (bottom). Dashed line: average unfolding force of wild-type P2. (**D**) Ring-knot occurrence (black) matches the exonuclease-resistance (red) in both wild type and mutant xrRNA.

The incidence of ring-knot formation in the FECs correlated remarkably well with the very different XR activities of the wild-type and mutant xrRNA (Fig. 4D). Notably, however, these results show that XR activity does not relate simply to high unfolding force: the variant pseudoknot formed by the mutant unfolded at ∼50 pN on average, not much lower than the ring-knot, yet apparently does not contribute to XR activity, else the mutant XR activity would be over 30% instead of only ∼9%. These results thus confirm that XR activity arises specifically from the mechanical resistance of the ring-knot fold.

By revealing the sequence of steps involved in the folding of the Zika xrRNA and the critical interactions leading to its mechanical resistance, these measurements show that the mechanical properties of the peculiar ring-knot topology of the xrRNA fold are essential to the ability of the viral RNA to evade degradation by cellular defenses. The intricate pattern of intermediates that must be followed to arrive at the resistant ring-knot structure suggests there may be several stages at which the ring-knot can be disrupted as a therapeutic intervention. Intriguingly, the central functional importance of the knot seen here contrasts starkly with the typical situation in knotted proteins, where knots—which occur much more commonly than in RNAs—generally play only an incidental role functionally (*27-29*). It will be interesting to see if this functional importance of mechanical resistance and the associated folding mechanism leading to resistant structures extend to other types of xrRNAs as they continue to be discovered.

## Acknowledgements

We thank Jeffrey Kieft for helpful discussions about xrRNAs, and Dustin Ritchie for assistance in designing the optical tweezers assay. This work was supported by the Canadian Institutes of Health Research, Alberta Innovates, and National Research Council Canada. The authors declare no competing interests.

## Supplementary Material

### Materials and Methods

#### Sample preparation

Constructs for SMFS measurements were generated similar to previous work (*21*). Briefly, the sequence for the wild-type Zika xrRNA (or C22G mutant) was inserted into a pcDNA5/FRT/TO plasmid between the HindIII and BamHI restriction sites (see Table S1). The resulting transcription template, which contained the xrRNA flanked by linker regions on either side, was then amplified by PCR and transcribed *in vitro* using T7 RNA polymerase. Two single-stranded (ss) DNA handles (one labeled with biotin and complementary to the 840 nt on the 3′ end of the transcript, the other labeled with digoxigenin and complementary to the 2,107 nt on the 5′ end of the transcript) were produced by asymmetric PCR from double-stranded DNA PCR products corresponding to the flanking handle sequences. The handles were thermally annealed with the RNA transcript, then incubated with 600-nm and 820-nm diameter polystyrene beads labeled respectively with avidin DN and antidigoxigenin, to create dumbbells. Dumbbells were placed in measuring buffer (50 mM MOPS pH 7.0, 130 mM KCl, 4 mM MgCl_2_, 50 U/mL Superase•In RNase inhibitor (Ambion)) containing oxygen-scavenging system (40 U/mL glucose oxidase, 185 U/mL catalase, and 8.3 mg/mL glucose) and inserted into a sealed sample chamber on a clean microscope slide in the optical trap. For measurements without Mg^2+^, the MgCl_2_ was removed from the measuring buffer and 1 mM EDTA added in its stead. For measurements using anti-sense oligos, 10 μM of the desired oligo was added to the measuring buffer.

#### SMFS measurements and analysis

FECs were measured with a custom-built, dual-beam optical trap similar to one described previously (*30*). Briefly, two orthogonally polarized beams from a 1,064-nm laser creating two traps were steered independently with acousto-optic deflectors. The motions of beads held in the traps were detected by collecting the light scattered by the beads onto position-sensitive diodes from two orthogonally-polarized 830-nm laser beams. The trap stiffnesses (0.54 and 0.56 pN/nm) were calibrated as described (*31*). Data were sampled at 20 kHz and filtered online at the Nyquist frequency with an 8-pole Bessel filter.

Contour length changes were determined from the FECs by fitting each state in the FEC to an extensible worm-like chain (WLC) model consisting of two WLCs in series: one for the duplex handles, the other for the unfolded ssRNA in each state. The extensible WLC was described by

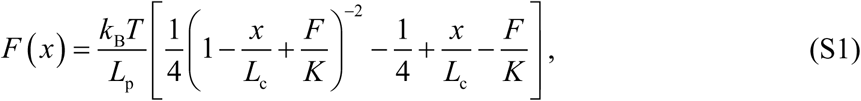

where *L*_p_ is the persistence length, *L*_c_ the contour length, and *K* the elastic modulus (*17*). The parameters *L*_p_, *L*_c_, and *K* for the duplex handles were first determined by fitting the natively-folded state (N) in the FECs. Then, the FECs for the various intermediate states and unfolded states were fitted by treating *L*_p_, *L*_c_, and *K* as fixed variables for the duplex handles (values determined from the previous fit) and *L*_p_ and *K* as fixed variables for ssRNA (values determined from the literature: *L*_p_ = 1 nm, *K* = 2,000 pN) (*32*). The contour length of unfolded ssRNA was therefore the only free parameter for fitting the intermediate and unfolded states. For those FECs that did not contain N, the unfolded state was instead used as the reference point for WLC fitting to measure the contour length of RNA unfolded in partially folded intermediates (see Tables S2-S4).

Structures were identified from FECs based primarily on the contour-length changes (Δ*L*_c_) found from WLC fits, which we compared to the values expected for the proposed structures. The total Δ*L*_c_ for completely unfolding a structure containing *n*_nt_ nucleotides is given by:

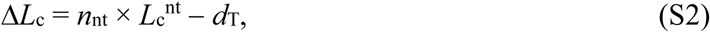

where *L*_c_^nt^ = 0.59 nm/nt is the contour length per nt (*33*), and *d*_*T*_ is the distance between the termini of the folded structure (as measured or estimated from the reported crystal structure). For example, natively folded Zika xrRNA has *d*_T_ = 3.0 nm and *n*_nt_ = 70, leading to Δ*L*_c_^total^ = 38.3 nm for complete unfolding. Individual helices were assumed to have *d*_T_ = 2.2 nm, the width of an A-form helix (*33*). Structural assignments were also informed by the unfolding and refolding forces observed. Unfolding and refolding of secondary structures alone, without any tertiary structures, typically occurs at forces in the range 10–25 pN, relatively close to equilibrium so that unfolding and refolding forces are similar (*26*). In contrast, tertiary structures typically unfold at forces > 25 pN, and usually display a wide distribution of unfolding forces and strong hysteresis when refolding (*18-22*). Only structural changes that alter the end-to-end length of the RNA can be observed directly by SMFS; conformational changes that leave the end-to-end length unchanged (e.g. breathing of base pairs or junctions) will remain undetected unless they also change the unfolding force sufficiently so that distinct unfolding force distributions can be identified.

**Fig. S1:**
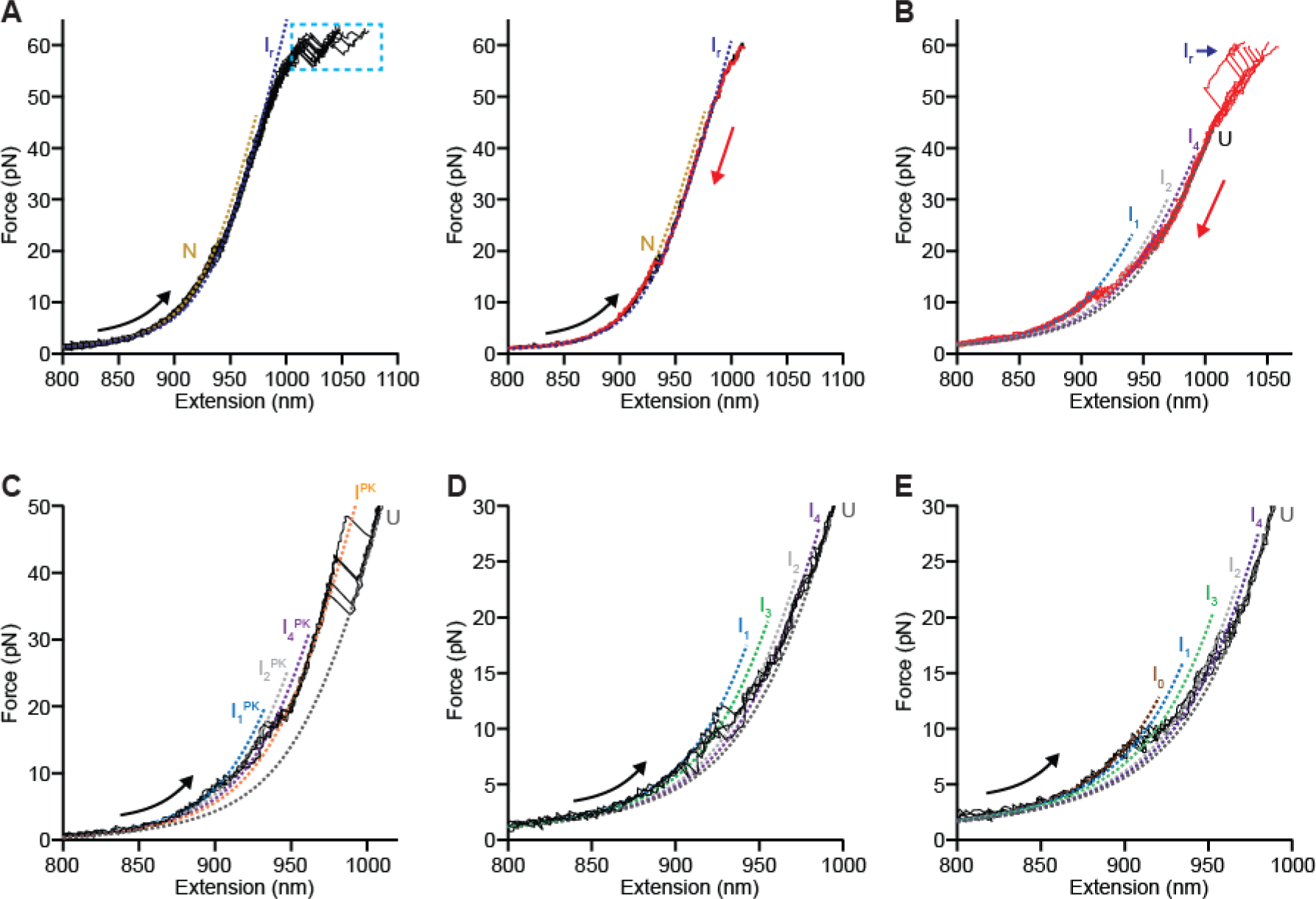
Representative FECs of wild-type Zika-xrRNA. (**A**) Multiple unfolding FECs (left panel) showing a mechanically resistant intermediate (I_r_) that remains after partial unfolding of the native state (N). I_r_ unfolding usually occurred during the overstretching transition at ∼60 pN (cyan), such that unfolding could not be distinguished from overstretching. If the unfolding force was not maintained at ∼60 pN for long enough during unfolding (right panel, black), then I_r_ typically stayed intact and returned to N in the subsequent refolding FEC (red). (**B**) In some FECs, I_r_ remained folded at the beginning of the refolding curve but subsequently unfolded to U while the force remained elevated. Refolding from U into I_1_ was then seen to occur through I_4_ and I_2_. (**C**) Representative unfolding FECs indicating a structure containing the pseudoknot without 5′-end threading, I_1_^PK^, which unfolds through distinct intermediates I_2_^PK^, I_4_^PK^ and I^PK^. (**D**) Representative FECs showing unfolding of native secondary structure only. (**E**) Representative FECs showing unfolding of secondary structure only but including an additional non-native helix in intermediate I_0_. Dotted curves show WLC fits to each state.

**Fig. S2:**
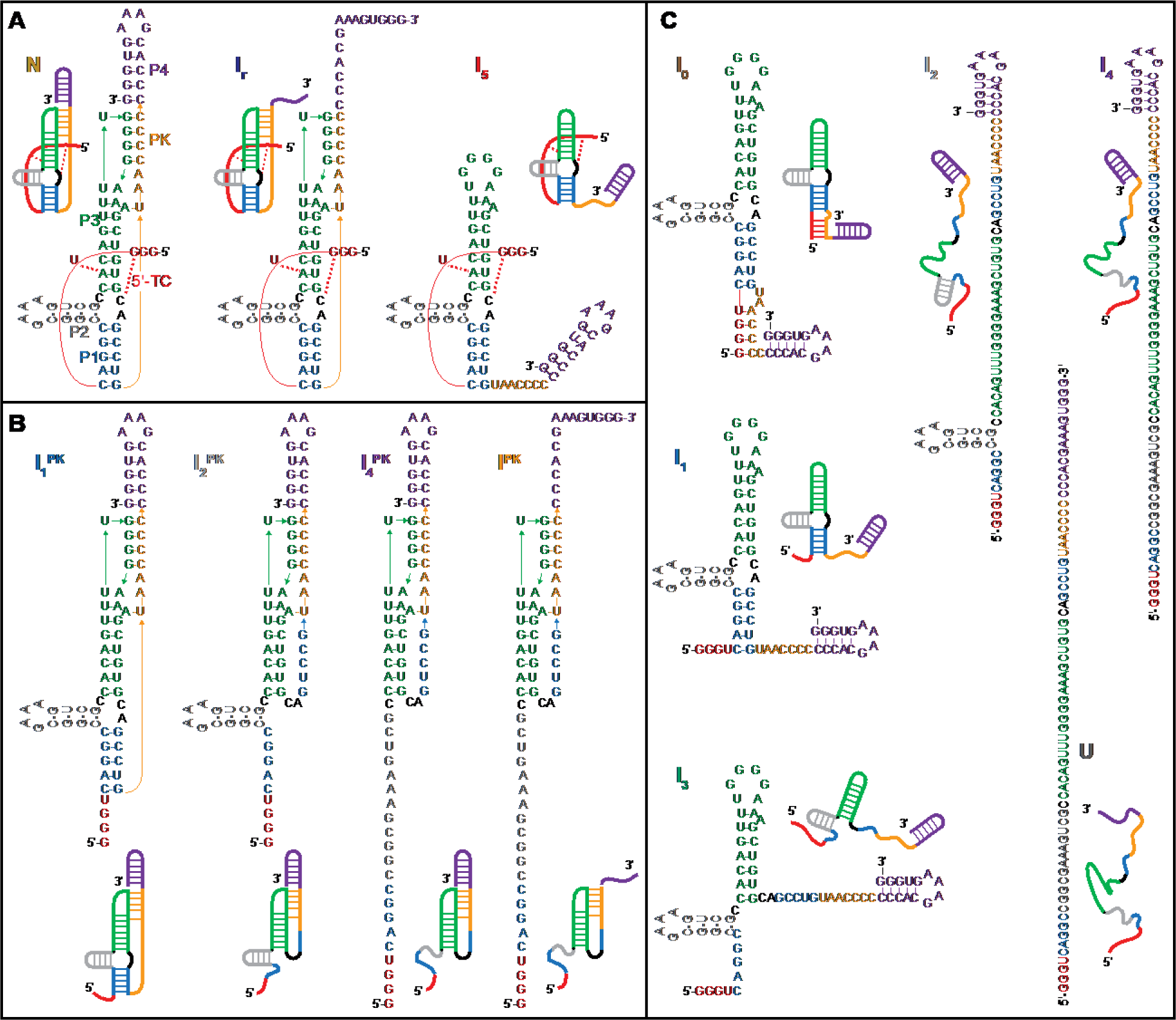
Structural models of states on unfolding and refolding pathways for wild-type Zika xrRNA under native conditions. (**A**) States with tertiary contacts stabilizing threading of the 5′ end. (**B**) States containing pseudoknots but no threading of the 5′ end. (**C**) States containing only secondary structure. The color coding is the same as in Fig. 1.

**Fig. S3:**
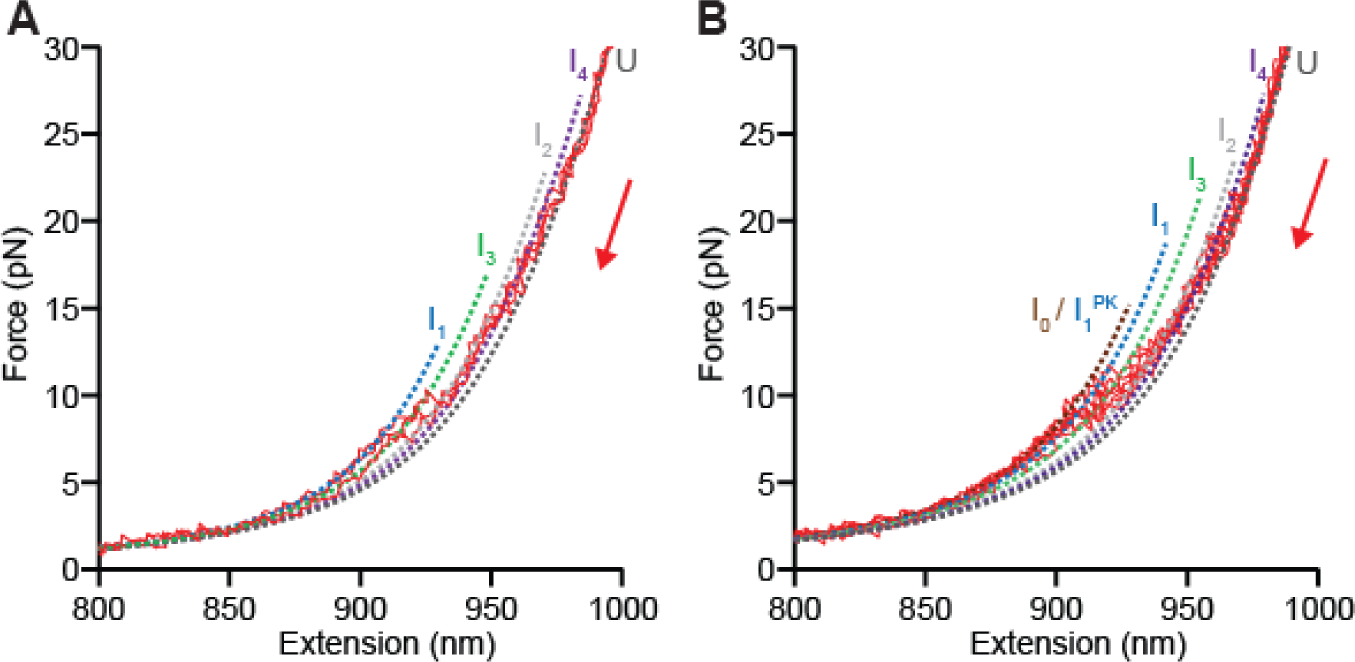
Refolding FECs. (**A**) Refolding FECs leading into state I_1_ show sequential formation of each helix. **(B)** Refolding FECs leading into I_0_ or I_1_^PK^ show a similar pattern, with an additional transition at low force. States I_0_ or I_1_^PK^ can only be distinguished in unfolding FECs by their unfolding forces, because their Δ*L*_c_ values are too similar (Table S2). Dotted curves show WLC fits to each state.

**Fig. S4:**
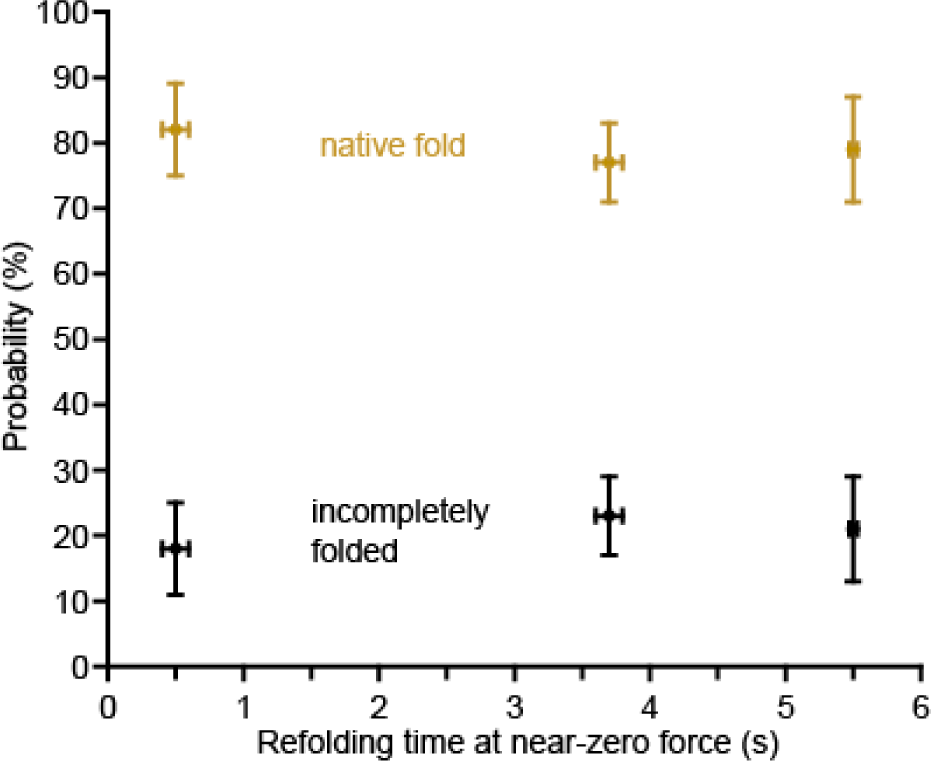
Effect of waiting time near zero force on xrRNA unfolding. The probability of finding the xrRNA in the native state (gold) as opposed to one of the incompletely folded states (black) is the same within error whether waiting 0.5 s (601 FECs from 17 molecules), 3.7 s (975 FECs from 35 molecules), or 5.5 s (1030 FECs from 19 molecules). Error bars represent s.e.m.

**Fig. S5:**
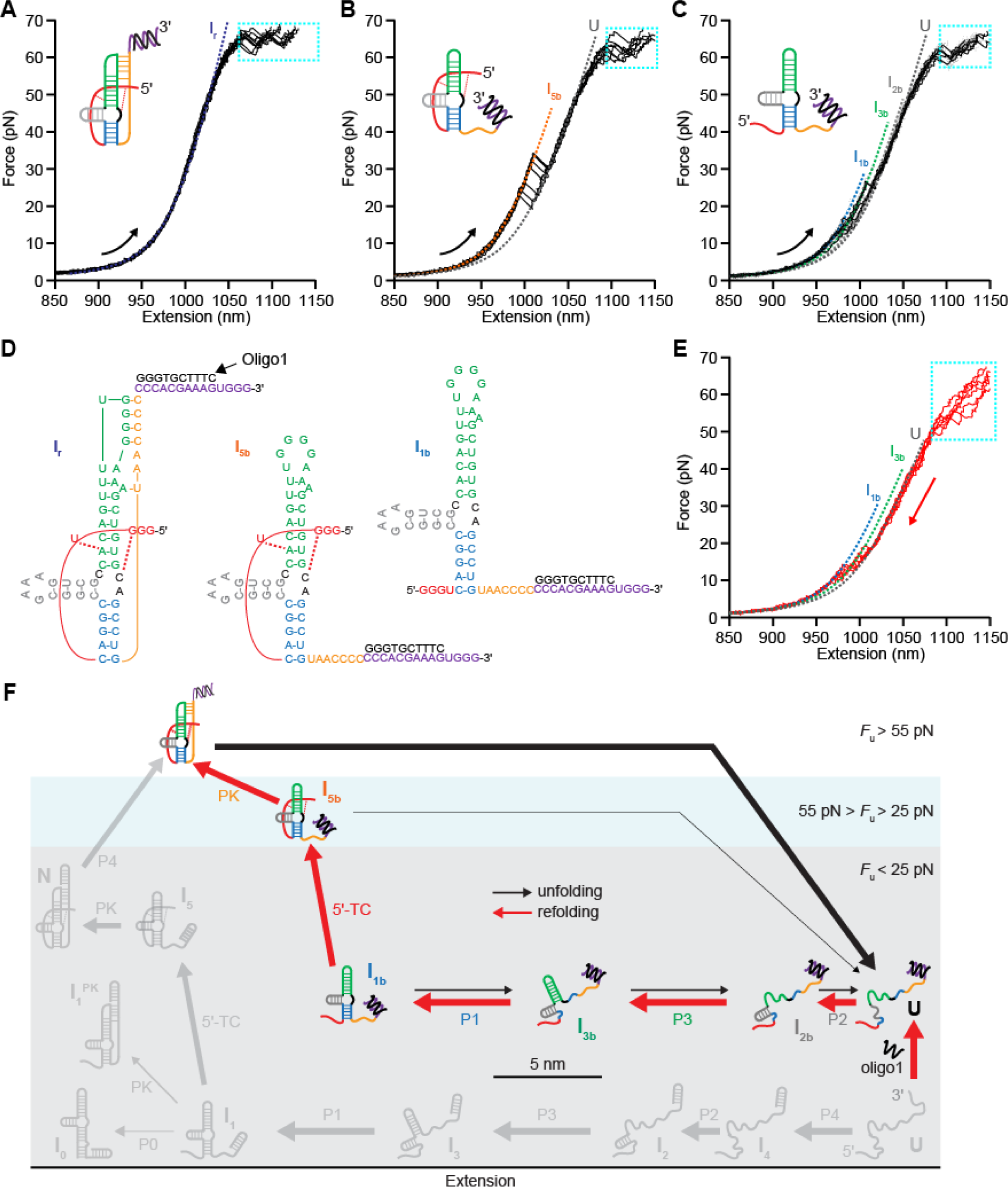
Effects of anti-sense oligo 1 on SMFS of Zika xrRNA. (**A**) Most unfolding FECs (90 ± 2%) show the extreme mechanical stability of I_r_ but no preceding ∼20-pN unfolding event, confirming that P4 unfolds prior to I_r_. Dashed lines show WLC fits to each state, cyan boxes show overstretching region, inset shows cartoon of structure before unfolding. (**B**) A very few FECs (∼0.3%) show unfolding forces in the range ∼25– 50 pN indicative of tertiary structure, and Δ*L*_c_ consistent with state I_5b_, where the 5′ end is threaded but the ring is not closed. (**C**) Some FECs (10% ± 2%) show unfolding of secondary structure only, consistent with state I_1b_. (**D**) Proposed structures of I_r_, I_5b_ and I_1b_ with oligo 1 bound. (**E**) Most refolding FECs show the formation of I_1b_; in some cases, transitions into I_r_ and/or I_5b_ are seen directly (*e.g*. in Fig 2B), in other cases, they occur at sufficiently low force to remain undetected. (**F**) Unfolding (black) and refolding (red) pathways in the presence of oligo 1. Arrow thicknesses indicate pathway probabilities. For comparison, the states and transitions prevented by binding of oligo 1 are shown in grey.

**Fig. S6:**
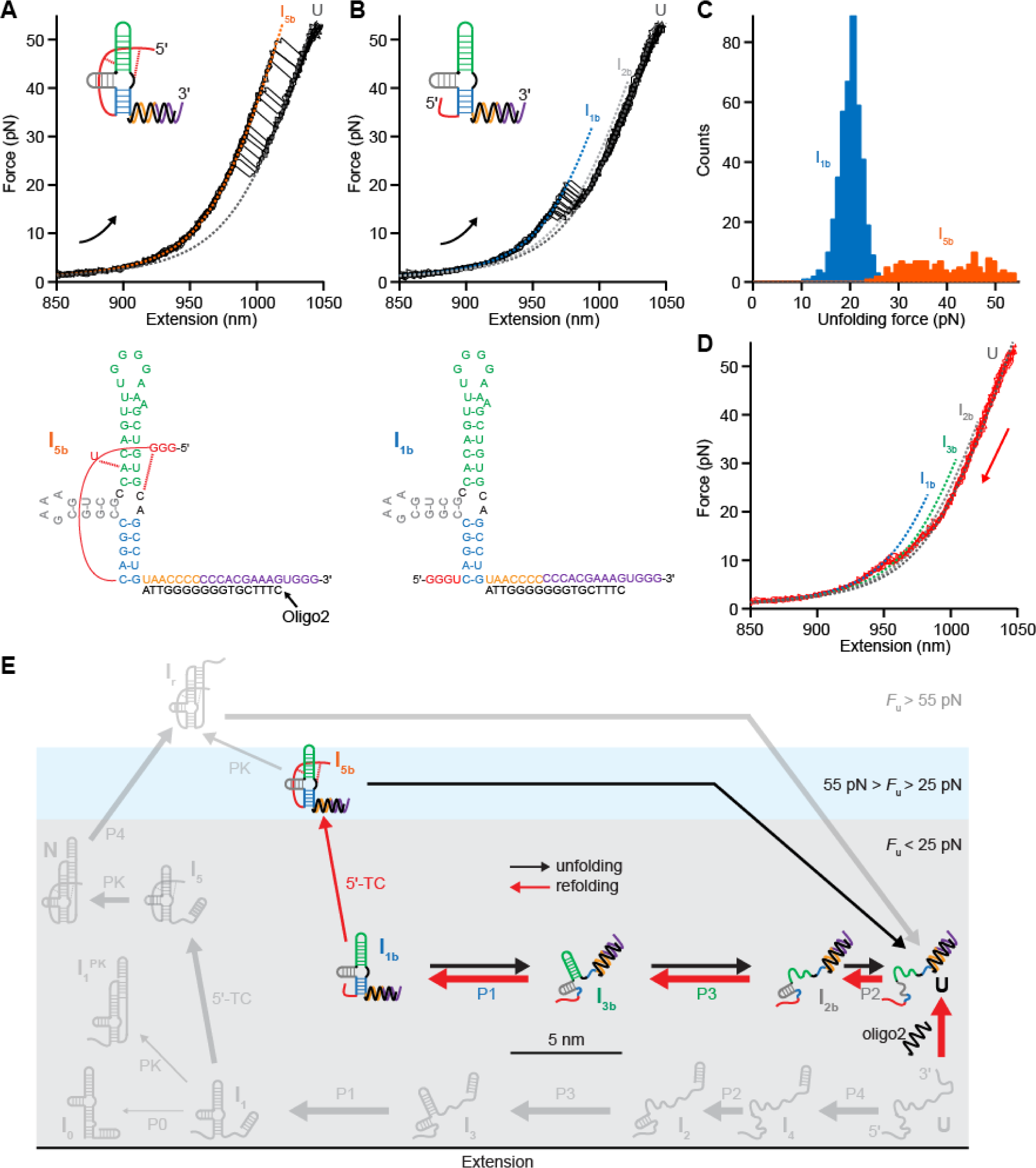
Effects of anti-sense oligo 2 on SMFS of Zika xrRNA. (**A**) Under half of FECs (41 ± 5%) show unfolding from I_5b_, with the 5′ end threaded but PK prevented by oligo binding. Dashed lines show WLC fits to different states, inset and cartoon show structure before unfolding with oligo 2 bound. (**B**) Over half of FECs (59 ± 5%) show unfolding from I_1b_. (**C**) States I_1b_ and I_5b_ can be distinguished by both their lengths (Table S3) and unfolding forces. (**D**) Most refolding FECs show the formation of I_1b_; in some cases, transitions into I_5b_ are seen directly (*e.g*. in Fig 2B), in other cases, they occur at sufficiently low force to remain undetected. (**E**) Unfolding (black) and refolding (red) pathways in the presence of oligo 2. Identification of I_5b_ indirectly proves the existence of the putative intermediate I_5_ (Fig. 3 and Fig. S2). Arrow thicknesses indicate pathway probabilities. For comparison, the states and transitions blocked by oligo 2 binding are shown in grey.

**Fig. S7:**
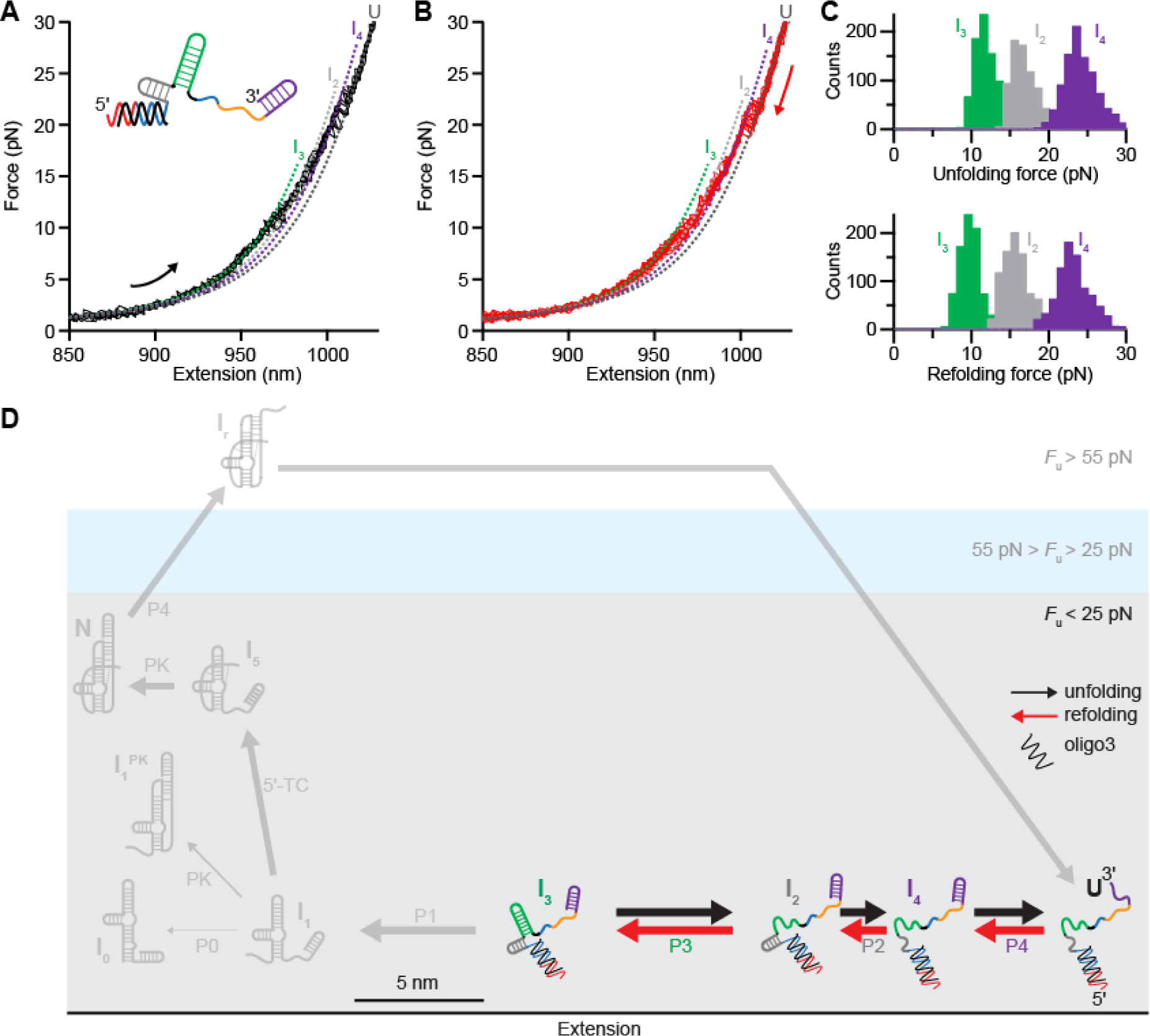
Effects of anti-sense oligo 3 on SMFS of Zika xrRNA. (**A, B**) All the unfolding (black) and refolding (red) FECs show the low-force transitions characteristic of secondary structure only, with length changes indicative of state I_3_. Dashed lines show WLC fits to different states, inset shows structure before unfolding with oligo 3 bound. (**C**) Unfolding (top panel) and refolding (bottom panel) force distributions show low hysteresis characteristic of helix unfolding/refolding. (**D**) Unfolding (black) and refolding (red) pathways in the presence of oligo 3. Arrow thicknesses indicate pathway probabilities. For comparison, the states and transitions blocked by oligo 3 binding are shown in grey.

**Fig. S8.**
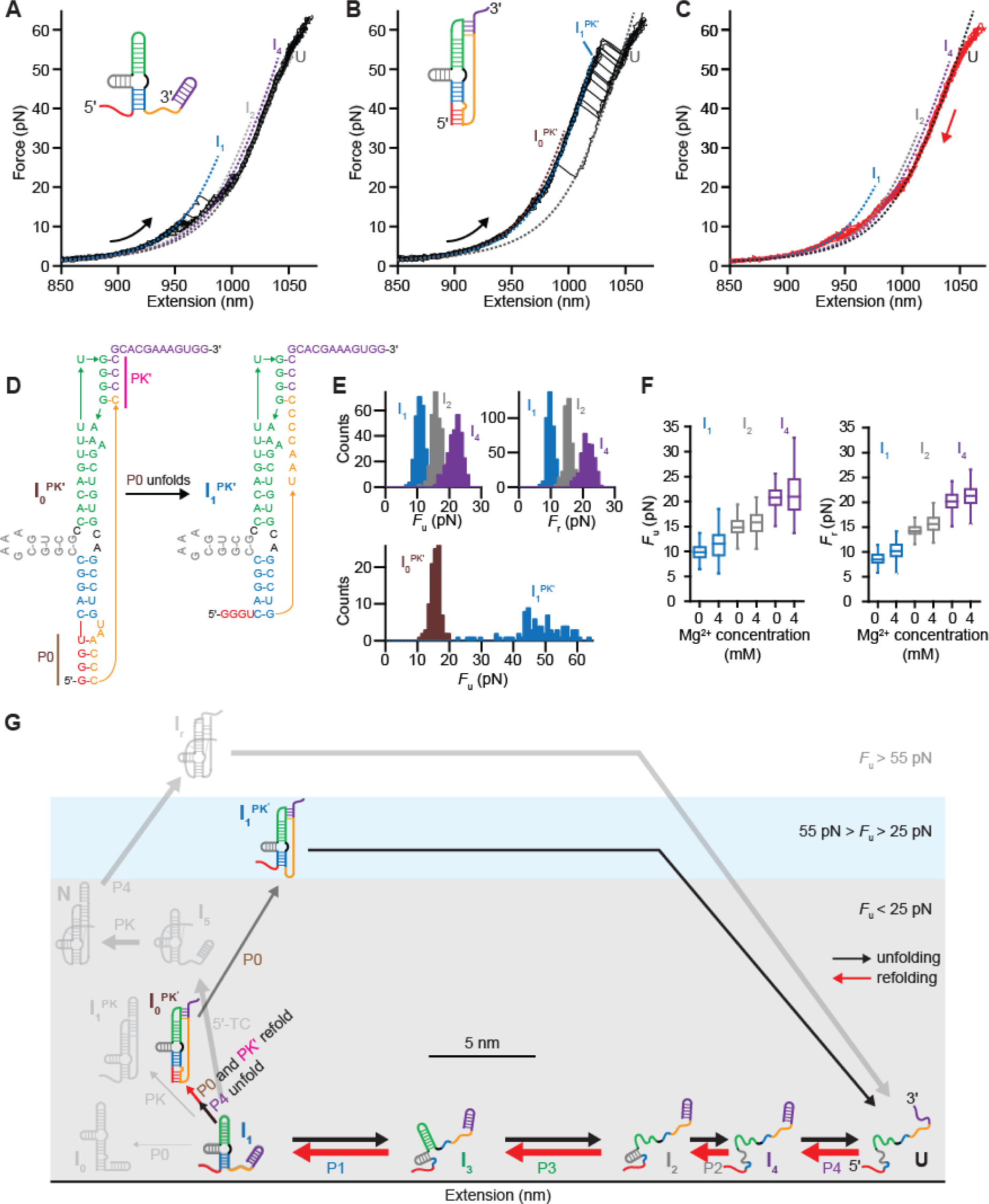
SMFS of Zika xrRNA in the absence of Mg^2+^. (**A**) Two-thirds of unfolding FECs (∼67%) show low-force transitions consistent with I_1_ unfolding (Fig. S2C). Dashed lines: WLC fits to each state. Inset: structure before unfolding. (**B**) The rest (∼33% of FECs) show high-force transitions consistent with state I_0_^PK′^ containing a non-native pseudoknot inhibiting P4 folding. (**C**) All refolding FECs show sequential folding of I_4_→I_2_→I_1_, indicating I_1_ is Mg^2+^-independent and suggesting I_0_^PK′^ derives from I_1_. (**D**) Cartoon of the I_0_^PK′^→I_1_^PK′^ transition. (**E**) Unfolding (*F*_u_) and refolding (*F*_r_) force distributions from FECs. High unfolding forces for I_1_^PK′^ indicate the presence of tertiary interactions. (**F**) Unfolding and refolding forces for I_4_, I_2_, and I_1_ show a small stabilization from Mg^2+^. (**G**) Unfolding (black) and refolding (red) pathways in the absence of Mg^2+^. Arrow thicknesses indicate pathway probabilities. For comparison, states and transitions dependent on Mg^2+^ are shown in grey.

**Fig. S9:**
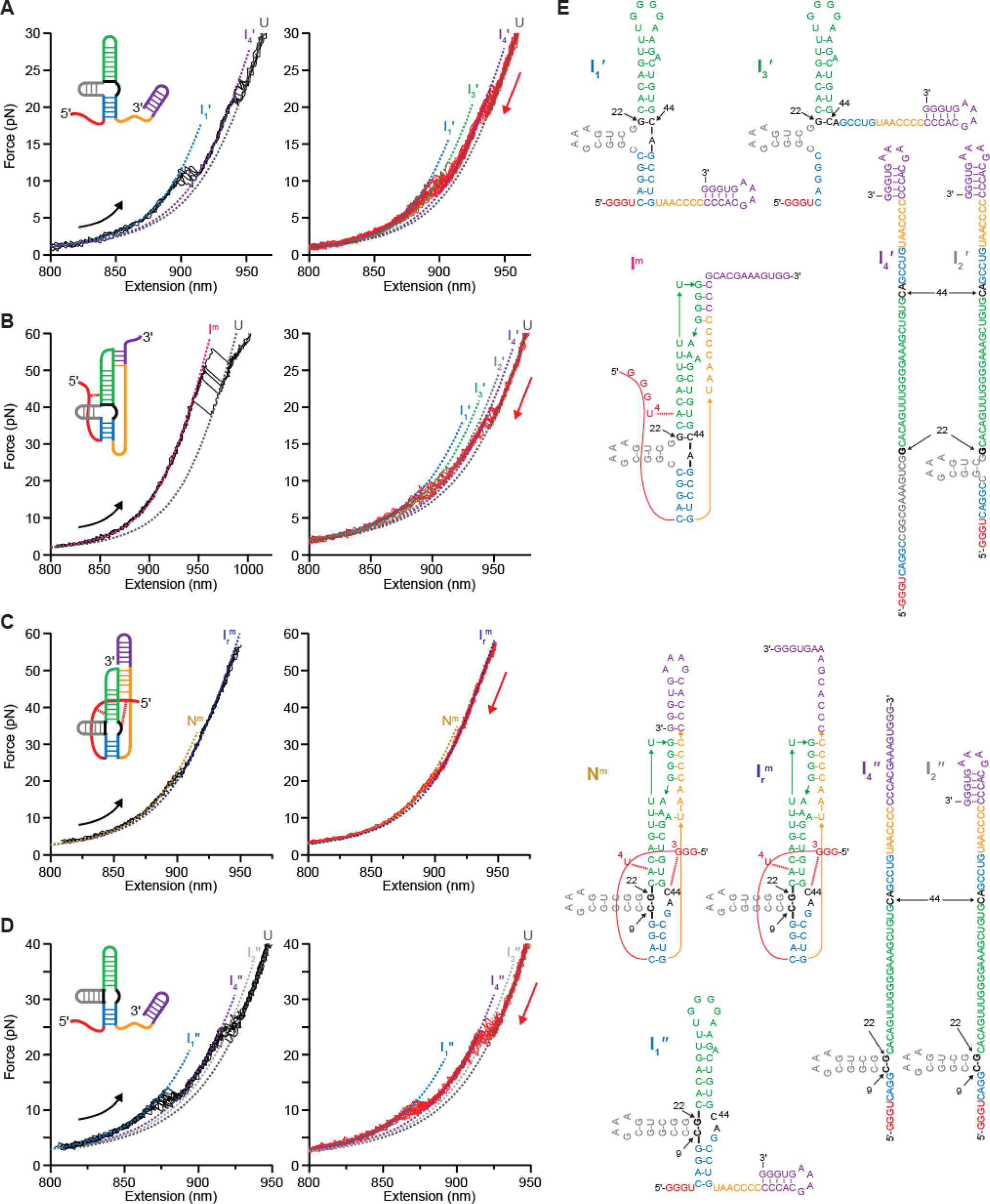
SMFS of the C22G mutant. (**A**) Roughly 15% of unfolding FECs (black) show low forces consistent with secondary structure only in state I_1′_, akin to I_1_ (Fig.S2C) but with shorter P2 and longer P3. Dashed lines show WLC fits to each state, inset shows structure before unfolding. (**B**) Roughly 17% of unfolding FECs (black) show a high-force state I^m^ consistent with partial formation of 5′-TC and the non-native pseudoknot PK′ but no 5′-end threading. Refolding FECs (red) show a pathway similar to I_1_ (Fig. S3A), although with shorter P2 and longer P3, indicating I^m^ derives from I_1′_. (**C**) Roughly 15% unfolding FECs (black) show the same extreme stability of the wild-type ring-knot, indicating a mutant ring-knot. (**D**) Roughly 53% unfolding (black) and refolding FECs (red) show low forces consistent with state I_1″_ (inset), akin to I_1_, containing secondary structure only but with lengthened P2. (**E**) Proposed structures of the states identified in (A-D). Basepairing of G22 with C44 prevents native 5′-TC but allows I_1′_–I_4′_ and I^m^ (with shorter P2 and longer P3) to form. Alternatively, base-pairing of G22 with C9 lengthens P2 but leaves C44 accessible for native 5′-TC formation, allowing formation of mutant states N^m^ and I_r_^m^ containing a ring-knot, in addition to intermediates I_1″_, I_2″_, and I_4″_.

**Fig. S10.**
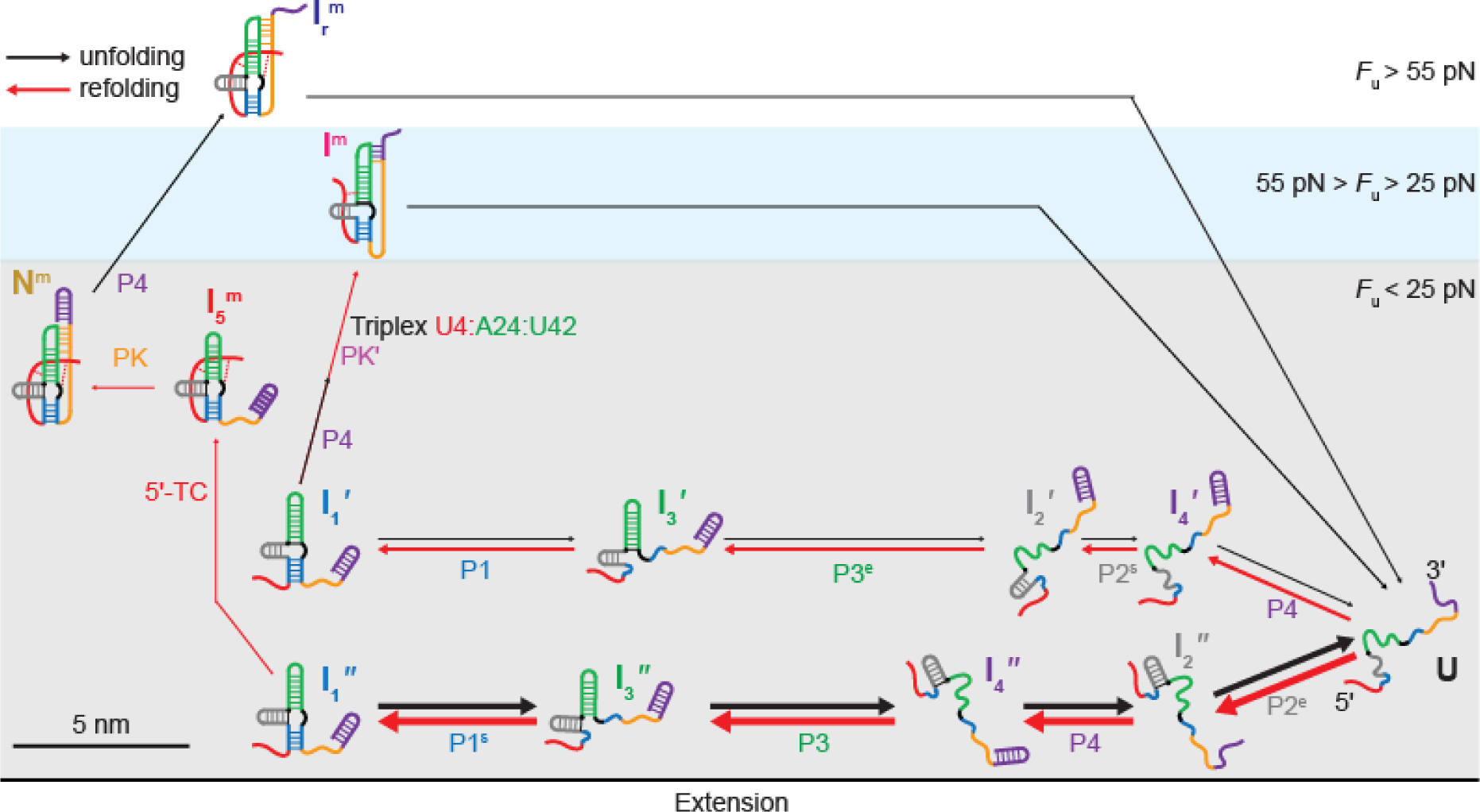
Unfolding and refolding pathways for C22G mutant. The two possible base-pairings for the mutated base (G22), with either C9 or C44, lead to heterogeneous pathways. Superscripts “e” and “s” denote respectively elongated and shortened versions of a helix. Arrow thicknesses indicate pathway probabilities.

**Table S1:**
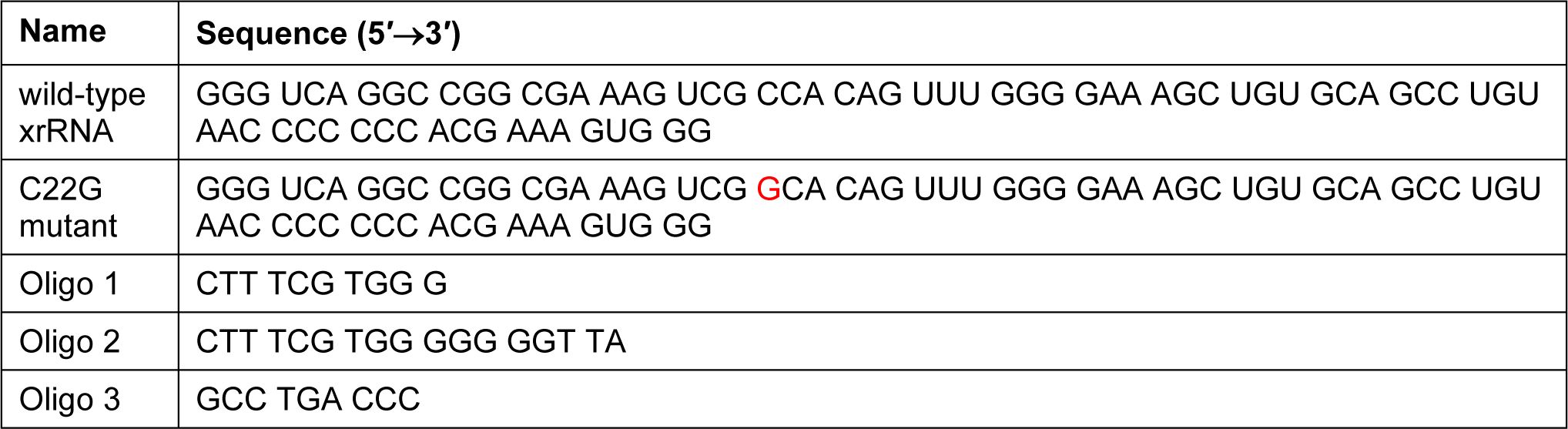
RNA and DNA sequences. Sequences of wild type Zika xrRNA, C22G mutant (mutation shown in red), and anti-sense DNA oligonucleotides used in SMFS measurements.

**Table S2:**
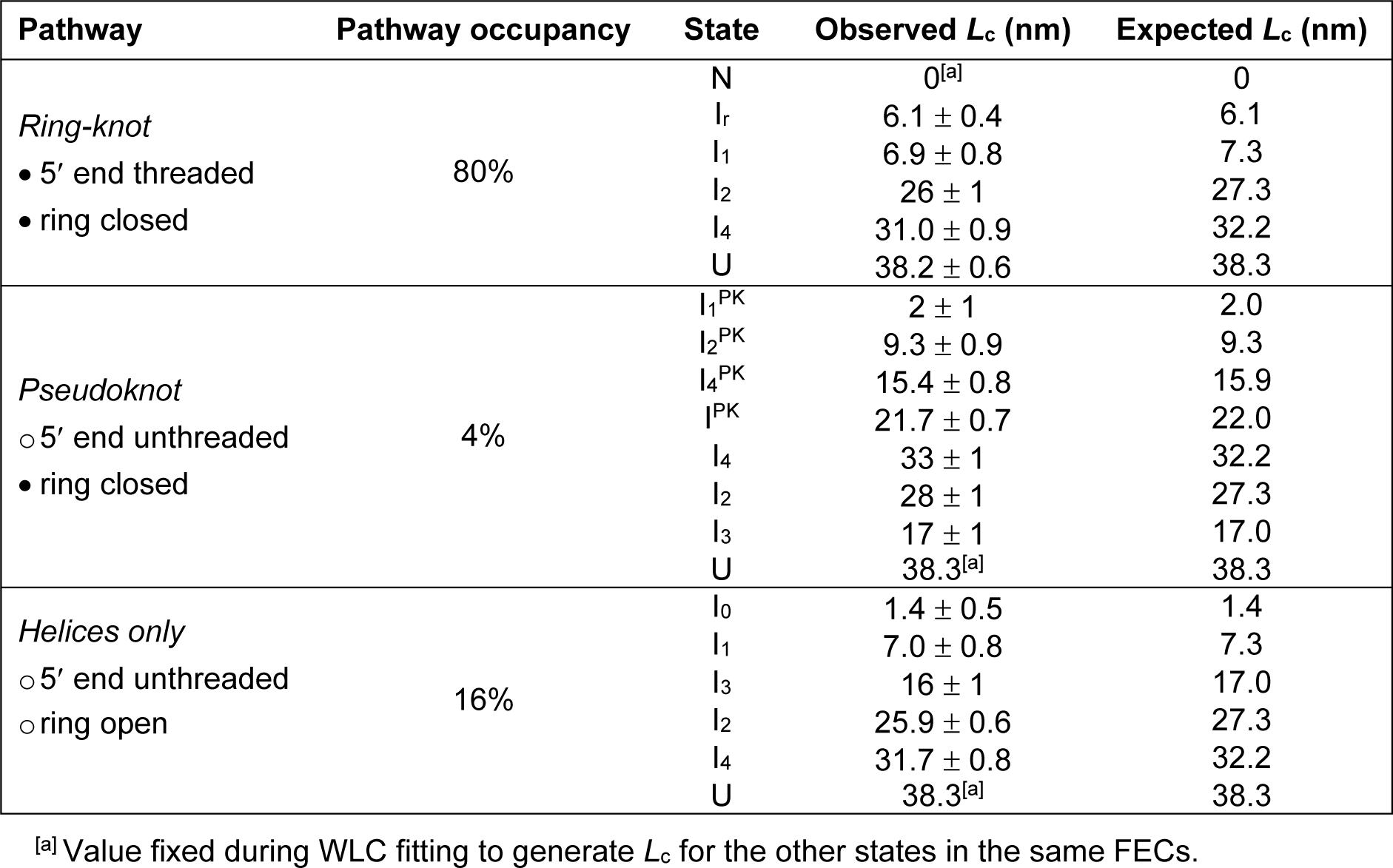
Contour lengths for wild-type Zika xrRNA unfolding and refolding. For each of the states identified in the unfolding and refolding FECs, from each of the three types of pathways observed (ring-knot, unthreaded pseudoknot, helices-only), the contour lengths relative to the native ring-knot, *L*_c_, are listed. The values observed (using a weighted average of the results from unfolding and refolding FECs) are compared to the values expected for the proposed structures based on the crystal structure. Errors in the observed contour length represent the standard error on the mean over all molecules measured (total of 5212 FECs from 69 molecules).

**Table S3:**
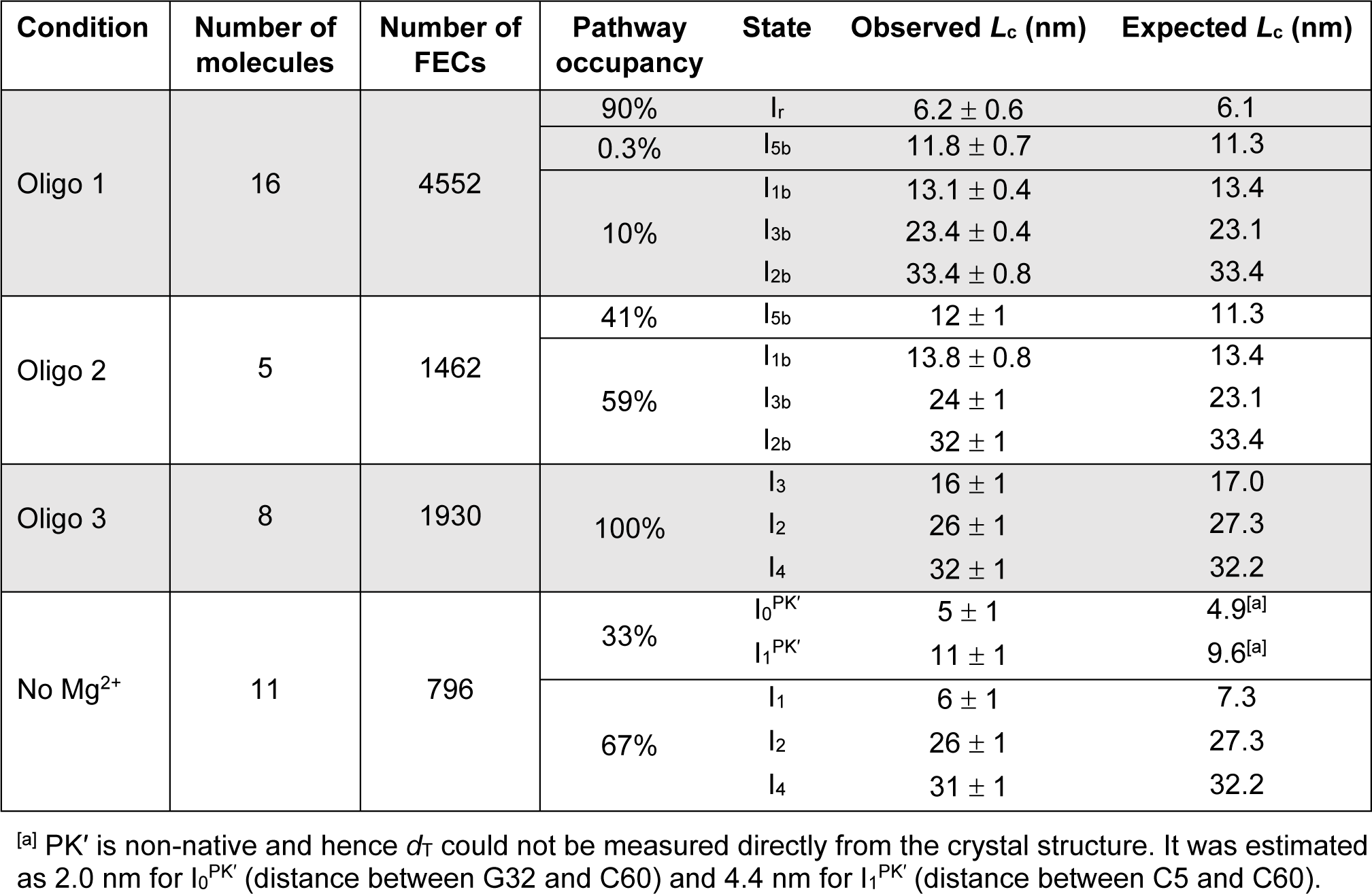
Contour lengths in presence of anti-sense oligos or absence of Mg^2+^. The observed and expected contour lengths relative to the native ring-knot are listed for each state identified on each pathway observed in each of four different conditions: the presence of one of the 3 anti-sense oligos, or the absence of Mg^2+^. Weighted averages of the results from unfolding and refolding FECs are compared to the values expected for the proposed structures. Errors represent s.e.m.

| Condition | Number of molecules | Number of FECs | Pathway occupancy | State | Observed $L_c$ (nm) | Expected $L_c$ (nm) |
| --- | --- | --- | --- | --- | --- | --- |
| Oligo 1 | 16 | 4552 | 90% | I<sub>r</sub> | 6.2 ± 0.6 | 6.1 |
|  |  |  | 0.3% | I<sub>5b</sub> | 11.8 ± 0.7 | 11.3 |
|  |  |  | 10% | I<sub>1b</sub> | 13.1 ± 0.4 | 13.4 |
|  |  |  |  | I<sub>3b</sub> | 23.4 ± 0.4 | 23.1 |
|  |  |  |  | I<sub>2b</sub> | 33.4 ± 0.8 | 33.4 |
| Oligo 2 | 5 | 1462 | 41% | I<sub>5b</sub> | 12 ± 1 | 11.3 |
|  |  |  | 59% | I<sub>1b</sub> | 13.8 ± 0.8 | 13.4 |
|  |  |  |  | I<sub>3b</sub> | 24 ± 1 | 23.1 |
|  |  |  |  | I<sub>2b</sub> | 32 ± 1 | 33.4 |
| Oligo 3 | 8 | 1930 | 100% | I<sub>3</sub> | 16 ± 1 | 17.0 |
|  |  |  |  | I<sub>2</sub> | 26 ± 1 | 27.3 |
|  |  |  |  | I<sub>4</sub> | 32 ± 1 | 32.2 |
| No Mg<sup>2+</sup> | 11 | 796 | 33% | I<sub>0</sub><sup>PK'</sup> | 5 ± 1 | 4.9<sup>[a]</sup> |
|  |  |  |  | I<sub>1</sub><sup>PK'</sup> | 11 ± 1 | 9.6<sup>[a]</sup> |
|  |  |  | 67% | I<sub>1</sub> | 6 ± 1 | 7.3 |
|  |  |  |  | I<sub>2</sub> | 26 ± 1 | 27.3 |
|  |  |  |  | I<sub>4</sub> | 31 ± 1 | 32.2 |

<sup>[a]</sup> PK' is non-native and hence $d_T$ could not be measured directly from the crystal structure. It was estimated as 2.0 nm for I<sub>0</sub><sup>PK'</sup> (distance between G32 and C60) and 4.4 nm for I<sub>1</sub><sup>PK'</sup> (distance between C5 and C60).

**Table S4:**
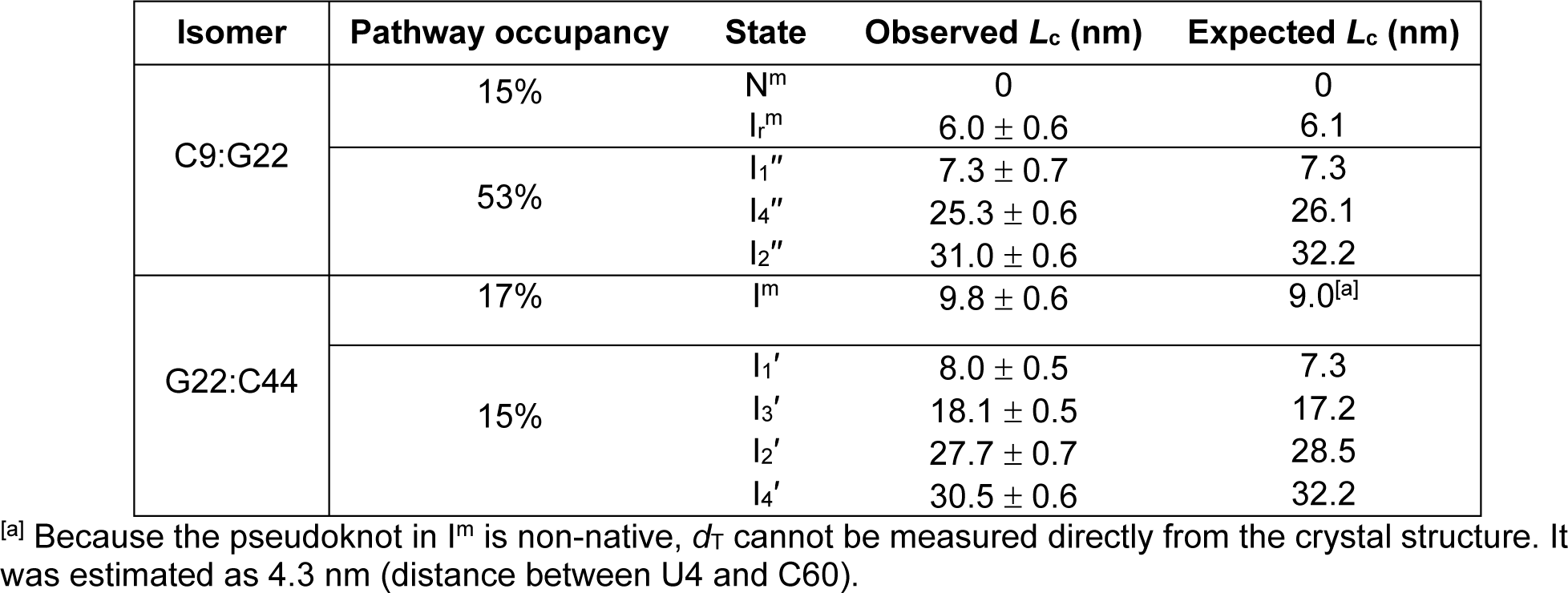
Contour lengths for C22G mutant. Contour length *L*_c_ (relative to N^m^) of the states identified in the unfolding and refolding FECs of the C22G mutant of Zika xrRNA, compared with the expected values. The observed and expected contour lengths relative to the state N^m^ are listed for each state identified on each pathway observed for the two isomers with different G22 base-pairing. Weighted averages of the results from unfolding and refolding FECs are compared to the values expected for the proposed structures. Errors represent s.e.m over all molecules measured (total of 1414 FECs from 11 molecules).

